# Polio Virotherapy of Malignant Glioma Engages the Tumor Myeloid Infiltrate and Induces Diffuse Microglia Activation

**DOI:** 10.1101/2022.04.19.488771

**Authors:** Yuanfan Yang, Michael C. Brown, Gao Zhang, Kevin Stevenson, Malte Mohme, Reb Kornahrens, Darell D. Bigner, David M. Ashley, Giselle Y. López, Matthias Gromeier

## Abstract

Malignant gliomas commandeer abundant inflammatory infiltrates with glioma-associated macrophages and microglia (GAMM) actively promoting tumor progression. Like all cells in the mononuclear phagocytic system, macrophages and microglia constitutively express the poliovirus receptor, CD155. Besides myeloid cells, CD155 is widely upregulated ectopically in the neoplastic compartment of malignant gliomas (and solid cancers in general). Intratumor treatment with the highly attenuated rhino:poliovirus chimera, PVSRIPO, yielded long-term survival with durable radiographic responses in patients with recurrent glioblastoma (Desjardins *et al*. New England Journal of Medicine, 2018). Here, we studied mechanisms of PVSRIPO immunotherapy in mouse brain tumor models to decipher contributions of myeloid vs. malignant cells to antitumor efficacy. PVSRIPO treatment caused intense engagement of the GAMM infiltrate associated with substantial, but transient tumor regression. This was accompanied by diffuse microglia activation and proliferation in the normal central nervous system (CNS) surrounding the tumor, extending to the ipsilateral and even the contralateral hemispheres. PVSRIPO-instigated microglia activation occurred against a backdrop of sustained innate antiviral inflammation, associated with induction of the PD-L1 immune checkpoint on GAMM. Combining PVSRIPO with PD1/PD-L1 blockade led to durable remissions. Our work implicates GAMM as active drivers of PVSRIPO-induced antitumor inflammation and reveals profound and widespread neuroinflammatory activation of the CNS-resident myeloid compartment by PVSRIPO.

## Introduction

Malignant gliomas comprise a diverse and heterogeneous group of primary CNS neoplasms that respond poorly to standard of care chemoradiation. They feature dense inflammatory infiltrates, shaping a tumor microenvironment (TME) populated with GAMM [35]. Microglia—derived from the embryonic yolk sac [19] and capable of self-renewal in the CNS [2, 27]—and CNS-associated macrophages are the principal innate immune cells in the CNS [40]. Malignant gliomas actively recruit and subvert these cells, ostensibly for tumor-accommodating pro-growth, angiogenic, and immune-skewing effects (reviewed in [39]).

Treatment of glioblastoma (GBM) with PVSRIPO achieved greater long-term survival vs. criteria-matched external controls with Overall Survival of 21% vs. 4% at 36 months [15]. Long-term survival after PVSRIPO was associated with objective radiographic responses with a median duration of >60 months. Polioviruses (PVs) use human CD155 (hCD155) as their sole host cell receptor [32]. Upon natural (oral) infection, PV targets certain lymphoid and intestinal epithelial cells and CD11c^+^ dendritic cells (DCs)/macrophages [44]. In chimpanzees, lymphatic structures are the principal sites of PV replication [5, 41]; widespread involvement of the lymphatic system was the defining systemic pathology of lethal PV infection in a large autopsy series mandated in a 1916 polio outbreak in the US [11]. Paralytic polio—the mechanisms of which remain obscure to this day—may be due to accidental viral targeting of anterior horn motor neurons expressing CD155 [12], possibly a neurodevelopmental remnant [25].

Wild type (wt) PV is highly cytotoxic with rampant viral growth in DCs/macrophages *in vivo* [44] and *in vitro* [45]. In contrast, PVSRIPO is engineered for profound attenuation [21], evident as absent neuropathogenicity after high-dose intracerebral (I.C.) inoculation in non-human primates [16] and in human subjects [15]. In DCs/macrophages, PVSRIPO exhibits a peculiar non-cytopathogenic phenotype, resulting in sublethal viral translation with lingering propagation driving sustained innate antiviral type-I interferon (IFN) dominant inflammation [9, 10, 36].

This scenario raises questions if PVSRIPO—analogous to wt PV tropism for DCs/macrophages *in vivo*—targets the glioma myeloid infiltrate. Moreover, are viral effects limited to the (injected) TME or do they extend to CNS-resident myeloid cells? In this work, we used I.C. immuno-competent murine tumor models for comprehensive histology, immunohistochemistry (IHC) and RNAseq studies of PVSRIPO immunotherapy. We report robust GAMM engagement and tumor regression, without evidence for direct viral damage to the neoplastic compartment. Beyond GAMM, we observed microglia activation immersing the tumor periphery and reaching deep into contralateral cortex. This was evident as microglia with increased cytoplasm and thickened ramified processes—highlighted on Transmembrane protein 119 (Tmem119) [4] and Ionized Ca^++^-binding adaptor protein 1 (Iba1) [1] staining—and profuse microglia proliferation upon PVSRIPO administration. RNAseq of the tumor region identified type-I/II IFN innate signaling gene signatures and upregulation of PD-L1 on GAMM. Drastic, but transient, antitumor effects of the acute neuroinflammatory response to PVSRIPO resulted in durable remissions upon combination with PD1:PD-L1 immune checkpoint blockade.

## Materials and methods

### Virus, cells, and transgenic mice

We utilized PVSRIPO produced as a good-laboratory-practice lot suitable for research purposes. Infectious cDNA [21] was digested with *Mlu*I, *in vitro* transcribed with T7 polymerase (Megascript, Thermo Fisher) and full-length viral RNA was transfected into HeLa cells with DMRIE-C in Opti-MEM (both Thermo Fisher) [17]. Progeny virus recovered from transfected HeLa cell cultures was quantified by plaque assay and amplified as described previously [17]. Upon manifestation of cytopathic effects, the infected cultures were harvested, subjected to 3 freeze thaw cycles, the lysate was centrifugated at 14,000g, and the resulting supernatant was filtered through 0.1 mM syringe filters (Pall) and subjected to centrifugation through a 100-kDa cutoff spin column (Millipore). We used CT2A^hCD155^ and B16^hCD155^ cells for I.C. implantation into C57Bl6 mice transgenic for human CD155 (hCD155-tg mice). Parental CT2A cells (a gift from Dr. P. Fecci, Duke University) and B16.F10 (ATCC) were subjected to lentiviral transduction with hCD155, followed by Fluorescence-Activated Cell sorting (FACs) with α-CD155-PE (BioLegend). Cells, confirmed mycoplasma negative (Duke Cell Culture Facility), were grown in DMEM (Invitrogen) with 10% fetal bovine serum (Sigma, #F0926) to 60-70% confluency and harvested for I.C. implantation. Homozygous hCD155-tg mice (a gift from S. Koike, Tokyo Metropolitan Institute of Medical Science, Japan) are maintained as a breeding colony. For survival studies, 8-12 wks old, ∼17-25g male and female mice were used, with roughly equal distribution in each treatment group; female mice were used for non-survival studies. Mice were housed in the Duke University Cancer Center Isolation facility under BSL2 conditions with 12-hr light/dark cycles, relative humidity of 50^±^20%, and temperature of 21^±^3°C.

### Intracranial Tumor Models

All animal procedures were performed under a Duke IACUC-approved vertebrate animal use protocol. Briefly, prior to surgery, hCD155-tg mice were shaved on the scalp and ear-tagged. During surgery, mice were under continuously inhaled isoflurane anesthesia and pain control (0.1 ml, 0.5% meloxicam S.C.). Mice were mounted onto stereotactic frames (Stoelting) when there was no toe pinch response; a heating pad (37^°^C) was used to maintain body temperature. The scalp was sterilized (betadine + 70% ethanol), and an incision was made by a #10 blade scalpel along the sagittal suture. The surgical field was washed with 3% H_2_O_2_ to expose the bregma, which was set as the origin (x=0, y=0, z=0) of the coordinate system. A 50 μl microsyringe (Hamilton, #80901) pre-filled with tumor cells homogenously suspended in 2.4% methylcellulose was fixed onto the frame, and a keyhole was created at (x=2.0mm, z=-0.5mm) by microdrill, so the 30Gy injecting needle could pierce through the skull freely, to a depth of 3.6 mm measuring from where the bevel tip intersects with the brain surface. Each mouse received 1×10^5^ CT2A^hCD155^ or 1×10^3^ B16^hCD155^ cells (5 μl), infused at 2.5 μl/min by an automatic injector. One min after completing the injection, the needle was slowly withdrawn, and the trepanation hole was sealed with bone wax (Covidien). The wound was closed with surgical glue (Vetclose, Henry Schein, Inc.) and mice were monitored during recovery until rolling over. PVSRIPO/mock treatment was performed on a day that is ∼1/3 of the median survival for each tumor model (day 6 for CT2A^hCD155^; day 5 for B16^hCD155^). Mice were divided into cohorts by a random number generator, and stereotactic intratumor infusion of PVSRIPO [5×10^7^ pfu in 5 μl DMEM)/mock (5 μl DMEM)] was performed using the keyhole created for tumor implantation. Following tumor implantation, mice were weighed every 2 or 3 days until reaching the humane endpoint, defined as >15% weight loss from the highest recorded weight, or until neurological symptoms (seizure, paralysis, etc.) were evident. Mice surviving long-term (3-times median survival, >60 days) were challenged with contralateral implantation of the same number of cells.

### Histology, Immunohistochemistry (IHC), Immunofluorescence (IF)

At the sample collection interval defined by the specific study, mice were euthanized, and brains were harvested for fixation in 10% formalin (Fisher Chemical) for 1-2 days and transferred to 70% ethanol for dehydration. Brain samples were cut coronally and paraffin embedded on the cut surface for sectioning. All histology slides in this study were coronal sections (7 μm for histology; 20 μm for RNA extraction) of the tumor implantation site using the needle track and the anterior commissure as anatomical landmarks. This ensured that all sections were at the center (and the largest cross section) of each tumor to allow for comparison among samples. Hematoxylin and eosin (H&E) staining followed a standard protocol at the Duke Pathology core facility. Sections were deparaffinized in xylene twice (10 min) and gradually rehydrated in 100%, 90%, 70% ethanol (5 min each). Sections were left in distilled water (5 min), dipped in hematoxylin (1 min), washed in tap water (5 min), dipped in Eosin Y (1% alcoholic; 0.5 min), gradually dehydrated twice in 95% (5 min each) and 100% ethanol (5 min each), stained with xylene twice (10 min), and mounted with Sub-X Mounting Medium (Leica). IHC was done on a Roche Ventana Discovery XT platform; formalin-fixed, paraffin-embedded (FFPE) sections were submitted for automated staining with Omni-map anti-Rabbit horseradish peroxidase-3,3’-diaminobenzidine (HRP-DAB). Slides were deparaffinized and subjected to IHC with primary antibodies diluted in 0.1% bovine serum albumin, followed by appropriate HRP-conjugated secondary antibodies (**Supplementary Table 1**), DAB staining, and hematoxylin and bluing agent counterstain. After staining, slides were washed in running soap water, dehydrated with a series of graded ethanol, cleaned with Sub-X, and mounted in Sub-X Mounting Medium (Leica). Manual cell counts were from three different fields selected ventral to the tumor, away from the peritumoral inflammation. IF was performed with a multiplexing approach (Visikol, Inc.). Slides were subjected to iterative rounds of staining with different antibodies (**Supplementary Table 1**), interspersed with antibody-stripping procedures. Slides were scanned and assembled for visualization with high resolution to allow for co-localization analyses. The technician acquiring the data was blinded to the treatments.

### RNA sequencing analyses

All samples used to collect bulk RNA stemmed from study series 3 (**Supplementary Table 3**). Two samples collected at day 6 (untreated) comprised the baseline; mock-treated samples from days 8, 12, 15 comprised the control groups. The mock day 12 group was complimented with one mock-treated day 10 sample (high similarity on principle component analysis). For the PVSRIPO-treated cohort, 3-4 samples were selected from study days 8, 12 and 15 (**Supplementary Table 3**). Bulk RNA was extracted with the RNeasy Plus Micro Kit (Qiagen) from the tumor region in 20 μM FFPE sections and submitted to Genewiz, Inc. for library preparation and sequencing. Fastq derived quality metrics were assessed via FastQC v0.11.8. Next, the paired-end reads were aligned to the GRCm38 (mm10) mouse reference genome via STAR v2.7.2b with 75 bp clipped from the 3’ ends to remove the Illumina adapter sequence. Subsequently, post-alignment quality metrics were assessed via the log file output of STAR as well as QualiMap v.2.2.1 to confirm genomic mapping quality. Quantification and generation of the raw counts matrix was performed via featureCounts v1.6.3. The raw counts matrix was then normalized, and DE was calculated via DeSeq2 R package v1.30.0. Genes with counts summed across all samples <10 were removed prior to the DE calculation. DE calculation was performed within RStudio v1.1.456 running R version 4.0.2. The normalized counts matrix was then entered into GSEA software version 4.0.3 and run with the Hallmark Gene Set Database with default parameter settings. From the GSEA results, the top 50 gene sets ranked by NES were run in a leading-edge analysis. Volcano plots were generated via the EnhancedVolcano R package v1.8.0; bulk RNA-Seq deconvolution was done via the mMCP-counter R package v1.0.0 using the log2 TPM normalized counts. The stacked bar plots were created using the ggplot2 R package v3.3.2. Viral load was assessed by alignment of the fastq files to the polio virus genome, NCBI Reference Sequence: NC_002058.3 using STAR.

### Quantification and statistical analysis

Survival data were analyzed by Kaplan Meier curves; when comparing two groups significance was determined by Log Rank test. For comparing histology measurements between treatment and control groups student *t* test with unequal variance was used; Cohen’s effect size of >0.8 was defined as a ‘large effect’. Other assay-specific statistical tests are indicated in the corresponding figure legends. P values of <0.05 were considered significant. All statistical analyses were performed using Prism 9 software version 9.0; error bars represent Standard Error of the Mean (SEM). All data points reflect individual specimens, independent experimental repeats, or mice.

## Results

### Optimizing mouse intracerebral tumor models for blinded neuropathologist review

CT2A is a 20-methylcholanthrene-induced murine anaplastic astrocytoma [43] that faithfully recapitulates key properties of human malignant glioma [30]. CT2A tumors have a high take rate, grow aggressively, and are resistant to immune checkpoint blockade [38]. Mouse CD155 does not function as a PV receptor; therefore, to recapitulate the clinical scenario, we transduced cells with hCD155 (CT2A^hCD155^) for implantation into hCD155-tg mice. To optimize I.C. mouse tumor models for histopathological analyses of all brains in all cohorts, we adjusted multiple variables to ensure consistent tumor growth contained within the right hemisphere. This included needle size/use of a drill for trepanation, inoculum (number of cells), and inoculation depth for maximal tumor take with minimal incidence of injection tract infiltration/extra-parenchymal growth. With these measures, we achieved 100% tumor take with 93.3% of tumors contained within the injected hemisphere per gross pathological exam (**Supplementary Table 2**).

### PVSRIPO therapy mediates early treatment effects in CT2A gliomas

Six days after I.C. CT2A^hCD155^ implantation in hCD155-tg mice, a single stereotactic infusion of PVSRIPO (5×10^7^ pfu) or mock (DMEM) was done (**Fig. 1**). This interval was chosen because histology of brains harboring CT2A^hCD155^ tumors 6 days post implantation revealed sizeable, established striatal growths (**Fig. 1**). In three independent executions of this experiment, tumor-bearing brains were harvested from mice euthanized at baseline (day 6), and from mice in the post PVSRIPO/mock cohorts collected at days 8, 10, 12, 15 or 18 (**Fig. 1**). Brains of all mice (n=55) enrolled in the 3 series were collected for histology (**Supplementary Table 3**). Because some mice reached humane end points (>15% weight loss, clinical symptoms) on/before day 18, the systematic part of this study (n=49) was not extended beyond day 15, to avoid attrition bias.

**Fig. 1.**
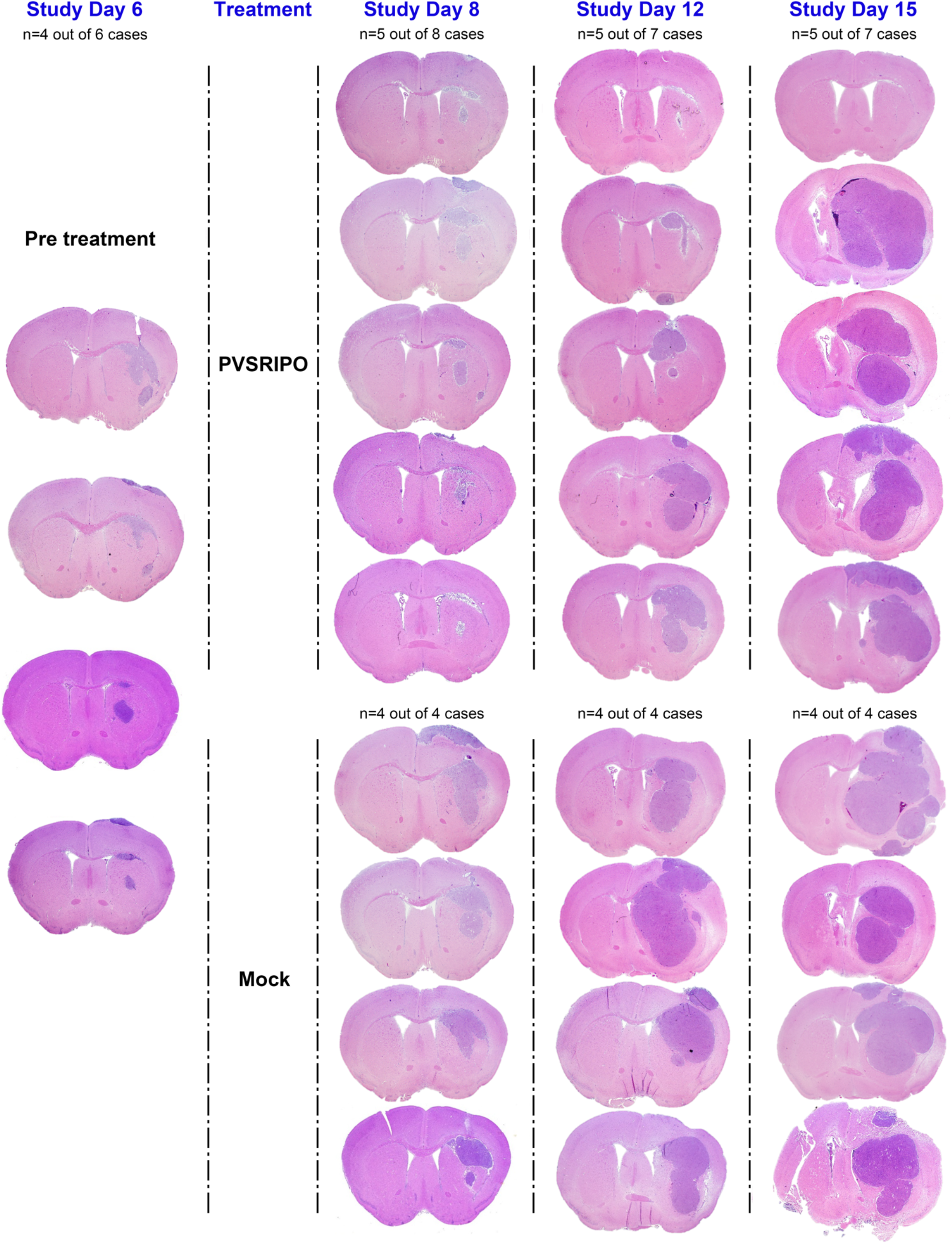
PVSRIPO therapy mediates a drastic, but transient antitumor effect. CT2A^hCD155^ cells implanted intracranially into the right hemisphere (day 0) gave rise to established gliomas on day 6 (baseline), at which time intratumor infusion of PVSRIPO (5×10^7^ pfu; in 5 μl DMEM) or mock (5 μl DMEM) was instituted. Tumor-bearing brains were collected at baseline (n=6), and post therapy (PVSRIPO or mock) at days 8 (n=12), 10 (n=9), 12 (n=11) and 15 (n=11) for histological analysis (see **Supplementary Table 3** for a listing of all samples). In each sample, coronal FFPE sections through the tumor implantation site were stained with H&E; tumors were evident as right hemisphere hypercellular masses. Typical micrographs showing the therapy response over time are displayed in the composite image, together with samples from baseline and matched controls.

Coronal FFPE sections of the tumor implantation site—using the needle track and anterior commissure as landmarks—were H&E stained for post-therapy histology in comparison with baseline and time-matched mock controls. Sample micrographs were assembled into a composite image to illustrate the treatment response of CT2A^CD155^ tumors to PVSRIPO over time (**Fig. 1**). This revealed a dramatic, but transient, antitumor effect evident as distinct histological appearance and reduced tumor size at day 8, followed by tumor relapse by day 15 (**Fig. 1**). At baseline (day 6), the average cross-sectional area of tumor was 1.2 mm^2^, which expanded by day 8 to 6.0 mm^2^ in the mock-, vs. 2.5 mm^2^ in the PVSRIPO arm (Cohen’s effect size >1.6; **Fig. 2c**). At day 15, the average cross-sectional areas of tumors were comparable in the mock (13.6 mm^2^) and PVSRIPO (14.2 mm^2^) arms (Cohen’s effect size 0.07; **Fig. 2c**). Occasionally, the early treatment response yielded complete pathological remission, leaving a scar at the implantation site (1/7 samples at day 15; **Supplementary Fig. 1**). No remissions were observed in the control cohort. Albeit rare, this is notable because CT2A glioma disseminates widely [30].

**Fig. 2.**
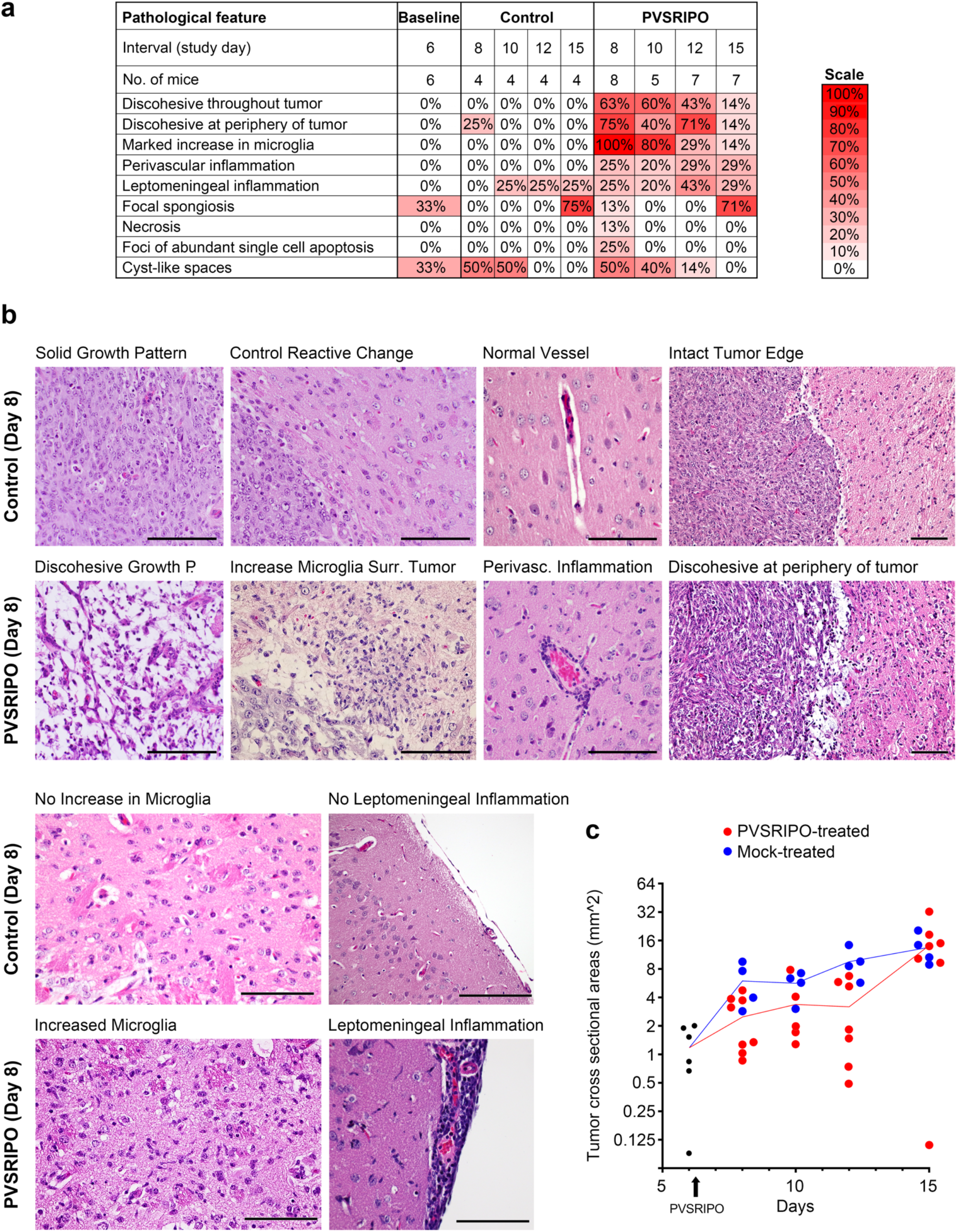
PVSRIPO treatment induces a transient histological response with sustained microglia inflammation. (**a**) To characterize the histologic features of PVSRIPO therapy in the CT2A^hCD155^ model, H&E-stained coronal FFPE sections at the tumor implantation site from each tumor-bearing brain collected at the indicated intervals, were submitted for histologic analysis by a board-certified pathologist (G.Y.L.) in a blinded fashion. Each slide was evaluated for the presence or absence of nine defined pathological features, and the results were summarized in color scales showing the prevalence of these features among all treated/control samples. (**b**) Certain histologic features were characteristic for PVSRIPO therapy, as shown in exemplary microscopic images compared to matched controls. Scale bars = 100 mm. (**c**) In each slide, the tumor cross sectional area (mm^2^) was calculated and plotted along the time course.

### The antitumor effect of PVSRIPO is histologically evident as marked microglia activation

To evaluate the PVSRIPO treatment effects noted in H&E-stained whole brain sections (**Fig. 1**), FFPE sections from the brains of 49 cases listed in Supplementary Table 3 were de-identified and submitted for blinded review by a Board-certified neuropathologist (G.Y.L.) to assess nine specific histological features (**Fig. 2**). This analysis revealed key features distinguishing PVSRIPO treated brains from the mock controls: (i) marked increase in microglia; (ii) discohesiveness throughout tumor; (iii) discohesiveness at periphery of tumor; and (iv) perivascular inflammation (**Fig. 2a, b**).

The predominant histological feature of the PVSRIPO response, present in all samples, was marked increase in microglia (**Fig. 2**). Microglia were evaluated in peritumoral brain parenchyma. Microglial activation was defined as aggregates of rod-shaped nuclei located immediately adjacent to the tumor. To meet criteria for an increased density of activated microglia, exclusion of other key cell types was required. The cells with rod-shaped nuclei had to lack visible cytoplasm. Abundant eosinophilic cytoplasm with ramified processes would support activated astrocytes, while abundant foamy cytoplasm would raise a differential of bone-marrow derived macrophages. Neither feature was present in regions identified as having activated microglia. While a mild increase in microglia (located mostly along the needle track) was found in mock-treated tumors relative to contralateral normal brain, we observed marked enrichment of activated microglia in the tumor periphery and throughout the brain after PVSRIPO treatment (see below).

### PVSRIPO causes diffuse microglia activation throughout the brain

Histopathological evidence for a microglial response to PVSRIPO (**Fig. 2**) requires rigorous determination of cell identity. Glioma implantation and virus inoculation induce CNS peripheral monocytic infiltrates which must be distinguished from tissue-resident microglia. Since Iba1 is also induced in activated CNS-associated macrophages or infiltrating peripheral monocytes [13], we added IHC for transmembrane protein 119 (Tmem119), which specifically marks microglia in the CNS [4], eg. in the context of RNA virus CNS infection [13].

To eliminate the tumor as a source for Iba1^+^ monocytic infiltration, we assessed microglia in non-tumor-bearing brains treated with PVSRIPO vs. mock (**Fig. 3**). IHC of Iba1 and Tmem119 demonstrated diffuse microglial activation, characterized as increased cytoplasm and thickened ramified processes, throughout the PVSRIPO-treated hemisphere (**Fig. 3a**). In mock-treated brains, Iba1^+^ cells occurred at the injection site only; these cells likely represent infiltrating monocytes, as they did not stain for Tmem119 in PVSRIPO/mock treated mice (**Fig. 3a**). CNS-wide distribution of Iba1^+^/Tmem119^+^ cells in PVSRIPO treated brains, excluding the injection site (**Fig. 3a**), provides definitive evidence for diffuse microglia activation by PVSRIPO. Higher magnification images of Iba1 (**Fig. 3b**) and Tmem119 (**Fig. 3c**) staining in cortex (distant from the injection site) revealed Tmem119^+^ cells with morphologic changes characteristic of activated microglia only in PVSRIPO treated brains (**Fig. 3c**).

**Fig. 3.**
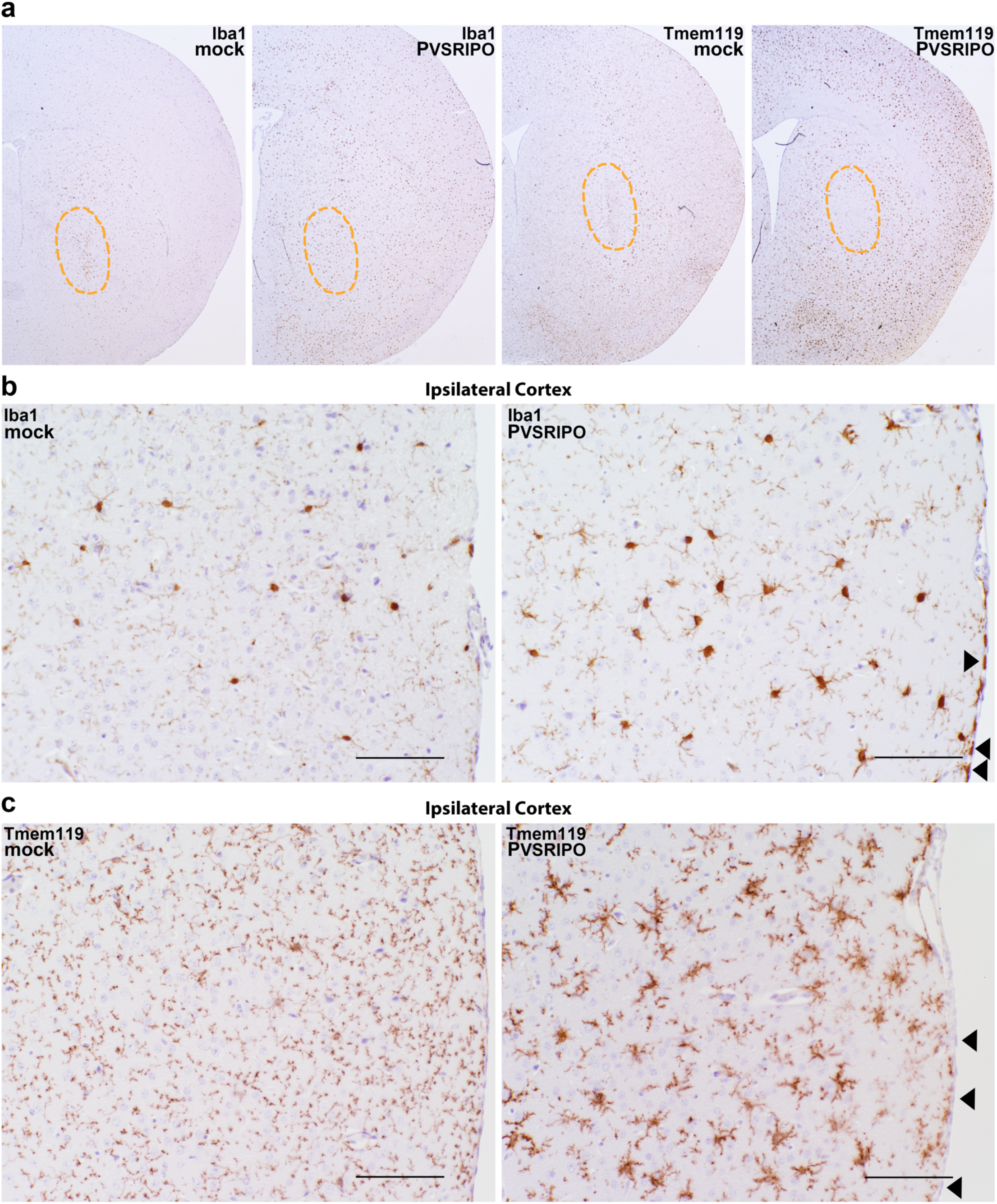
PVSRIPO therapy induces diffuse hemispheric microglia activation. Coronal FFPE sections of mouse brains treated with either PVSRIPO or a mock injection were stained with antibodies against (**a, b**) Iba1 and (**a, c**) Tmem119. (**a**) The injection site is marked by a dotted outline. Whole mount images of the injected hemisphere of mock-treated brains show Iba1^+^ signal only in cells clustering in the area of the needle track (left panel); these clusters did not stain with Tmem119 antibody (right middle panel). In PVSRIPO-treated brains, there was diffuse hemispheric Iba1^+^ and Tmem119^+^ signal; the region of the needle track was devoid of Tmem119^+^ cells (right panel). At higher magnification of an ipsilateral cortical area, Iba1 (**b**) and Tmem119 (**c**) marked microglia with increased cytoplasm and broad, thickened ramified processes in PVSRIPO-treated brains (right panels). Note Iba1^+^ (**b**; arrowheads), Tmem119^-^ (**c**; arrowheads) meningeal macrophages in PVSRIPO-treated brains. A microglial reaction and Iba1^+^ meningeal macrophages were absent in mock-treated brains (left panels).

### Confirmation of antitumor effects of PVSRIPO in an intracerebral B16 model

To corroborate our findings in CT2A^hCD155^ murine glioma, we selected the I.C. B16.F10 (melanoma) model transduced with hCD155 [10]. B16 is spontaneous and one of the least intrinsically immunogenic mouse tumor models [28]. PVSRIPO/mock treatment of I.C. B16^hCD155^-harboring mice was done on day 5 (instead of day 6 with CT2A^hCD155^) and brains were harvested on day 7 (instead of day 8 with CT2A^hCD155^) due to shorter median survival (∼14.5 days for B16^hCD155^ vs. ∼19.5 days for CT2A^hCD155^). Blinded pathological review of mock (n=2) vs. PVSRIPO-treated (n=3) B16^hCD155^-bearing brains (study day 7), revealed similar treatment effects: mock-treated tumors had clearly delineated borders with minimal inflammation, whereas PVSRIPO-treated tumors were smaller, discohesive, admixed with inflammation and markedly increased GAMM in and around the tumor with prominent peritumoral and perivascular inflammation (**Supplementary Fig. 2a, b**). The day 8 post-PVSRIPO antitumor effect noted in CT2A^hCD155^ was evident in Ki67 IHC (**Supplementary Fig. 2c**) and CD68 IHC signal was profusely enhanced in the tumor bed and surrounding brain (**Supplementary Fig. 2d**).

### PVSRIPO engages GAMM and causes widespread microglia activation and proliferation in the CNS

Having confirmed the microglial nature of Iba1^+^ cells throughout the ipsilateral hemisphere by presence of both highly ramified morphology and Tmem119 positivity (**Fig. 3**), we selected representative slides from PVSRIPO/mock-treated CT2A^hCD155^ glioma-bearing mice collected on day 8 (n=2 mock/4 PVSRIPO) for multiplex IF staining of Iba1, Ki67, PD-L1 (**Fig. 4a, b**) and confirmatory IHC of Iba1 (**Fig. 4c, d**). We focused on three regions: a central portion of the tumor, ipsilateral tumor-adjacent normal brain close to the anterior commissure (**Fig. 4a-d**), and distant contralateral cortex (**Fig. 4c, d**). Iba1 staining delineated tissue-resident microglia with characteristic ramified processes that emanate from a cell body in peritumoral normal brain (**Fig. 4a-d**). In PVSRIPO-treated brains, the presence of activated microglia was pervasive, with prominent microglia activation, both in peritumoral normal brain and in contralateral cortex (**Fig. 4b, d**). Profuse microglia activation was evident in all virus-treated brains, but none of the mock-treated controls. We confirmed microglia identity by staining with Tmem119 in adjacent sections (data not shown). Further signs of engaged microglia in PVSRIPO-treated brains were evident upon multiplex staining with Iba1 and Ki67 (**Fig. 4a, b**). In mock-treated animals, ipsilateral normal brain was devoid of Ki67 staining (**Fig. 4a**), while in PVSRIPO-treated animals Iba1^+^ cells with highly ramified processes co-stained with Ki67 (**Fig. 4b**), supporting widespread active microglia proliferation in brain parenchyma.

**Fig. 4.**
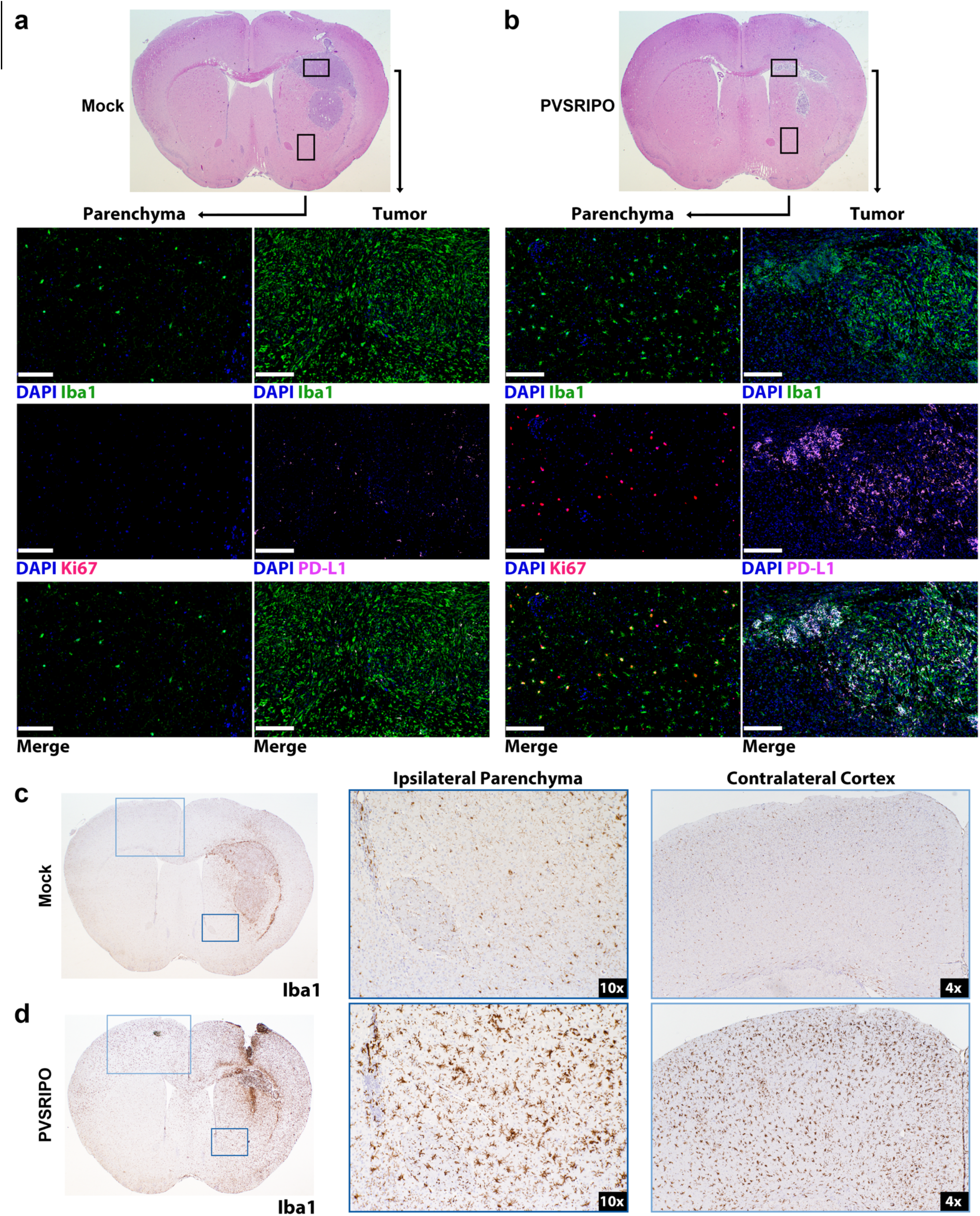
PVSRIPO therapy induced proinflammatory GAMM engagement and widespread activation and proliferation of tissue-resident microglia in the CNS. (**a, b**) Representative cases from day 8 post mock (**a**) or PVSRIPO (**b**) (see Figure 1) were analyzed by IF as shown. Areas representing the malignant lesion and an area in the adjacent brain parenchyma are indicated by boxes. Staining with DAPI (nuclear stain), Iba1, Ki67 (proliferation marker), PD-L1 of both regions in isolation or merged is shown as indicated. Scale bars = 100 μm. (**c, d**) Representative cases from day 8 post mock (**c**) or PVSRIPO (**d**) were analyzed for Iba1 IHC for documentation of microglia morphology. Two areas studied at higher magnification in the ipsilateral tumor-adjacent normal brain (dark blue frame), or the contralateral normal cortex (light blue frame) are indicated, highlighting widespread microglia activation with increased cytoplasm and thickened processes in the PVSRIPO-treated brains (**b, d**) compared to mock-treated brains (**a, c**).

Beyond microglia activation in ipsilateral/contralateral brain, PVSRIPO induced profound proinflammatory engagement in the TME. In mock-treated animals, PD-L1 staining in the tumor proper was sparse (**Fig. 4a**). In contrast, abundant PD-L1 co-staining with Iba1 in the tumor bed in PVSRIPO-treated mice indicated type-I IFN mediated proinflammatory activation of GAMM (PD-L1 is an IFN-stimulated gene; PVSRIPO induces PD-L1 in human DCs *in vitro* [36]; **Fig. 4b**). Iba1^+^ cell activation extending to normal ipsilateral brain and the contralateral hemisphere (**Supplementary Fig. 3b**), and active microglia proliferation in normal brain parenchyma (**Supplementary Fig. 3f**) were also observed in all cases of the I.C. B16^hCD155^ model treated with PVSRIPO that were submitted for multiplex IF analysis (n=3).

To further document the PVSRIPO treatment effect, we co-stained sections from the brains of CT2A^hCD155^-bearing mice after PVSRIPO/mock treatment by multiplex IF staining of CD45, PD-L1 and Ki67 (**Fig. 5**). In mock-treated animals, the CD45^+^ compartment remained well organized, interwoven with and mostly restricted to the neoplastic lesion itself (**Fig. 5a, b**) with scarce and punctate PD-L1 expression (**Fig. 5c**). Accordingly, Ki67 staining was intense in the neoplastic lesion, with some scattered signal in the peritumoral periphery (**Fig. 5d**). In PVSRIPO-treated animals, however, CD45 staining was no longer confined to the malignant lesion, but extended far into brain parenchyma, forming a vast, undulating inflammatory reaction that engulfed much of the tumor-bearing hemisphere (**Fig. 5e, f**), replete with PD-L1 signal (**Fig. 5g**). Ki67^+^ cells were greatly diminished in the tumor itself (in accordance with reduced tumor size; **Fig. 2c**) and were present throughout the hemisphere (**Fig. 5h**). In PVSRIPO-treated animals, Ki67^+^ cells located outside the tumor region largely co-stained with Iba1 and demonstrated highly ramified processes, suggesting proliferating microglia (**Fig. 5f**; **Supplementary Fig. 3f**); or, Ki67^+^ cells co-stained with CD3 in a tight perinuclear pattern, suggesting proliferating T cells (**Supplementary Fig. 5b**). Calculation of the density of CD45^+^ cells in CT2A^hCD155^ tumor-adjacent normal brain demonstrated that PVSRIPO treatment induced an ∼10-fold influx of such cells into peritumoral parenchyma; the same effect was observed in the B16^hCD155^ model (**Supplementary Fig. 4a**). Automated quantification of CD45, Iba1, PD-L1, CD3 and Ki67 signal in CT2A^hCD155^ tumor and in ipsilateral brain of mice treated with mock vs. PVSRIPO are shown in Supplementary Figure 4b, c.

**Fig. 5.**
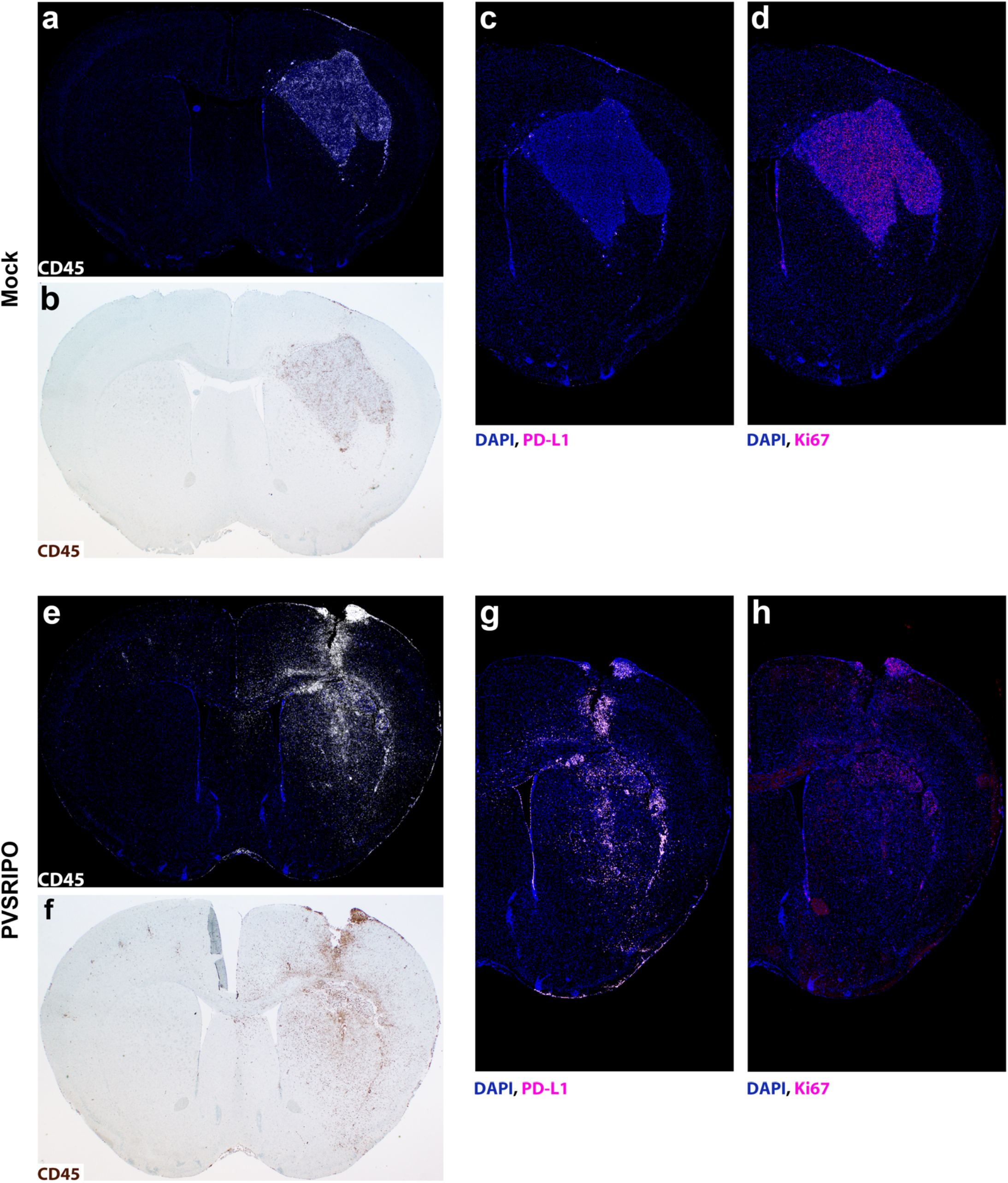
PVSRIPO immunotherapy is associated with profound inflammatory responses and upregulation of PD-L1 in the tumor microenvironment, involving the entire tumor-bearing hemisphere. Mock-(**a**-**d**) and PVSRIPO-treated (**e**-**h**) CT2A^hCD155^ tumors (day 8) were stained for CD45, Ki-67 and PD-L1 expression as shown.

### Increased intra- and extra-tumoral T cell density and proliferation after PVSRIPO

Co-staining with CD3 suggested that some of the Ki67^+^ cells in PVSRIPO/mock-treated CT2A-bearing brains were T lymphocytes (**Supplementary Fig. 5**). However, in contrast to mock-treated mice, where Ki67^+^ CD3^+^ cells occurred mostly within the tumor bed (**Supplementary Fig. 5a**), such cells were disseminated throughout the peritumoral circumference of PVSRIPO-treated CT2A^hCD155^ (**Supplementary Fig. 5b**) and B16^hCD155^-bearing mice (**Supplementary Fig. 6**). In aggregate, our IF and IHC studies revealed a broad range of proinflammatory effects engulfing the CNS after intratumor PVSRIPO treatment. The effects were noted for their quality, eg. profuse microglia proliferation in the normal CNS and robust PD-L1 induction on GAMM and in the tumor-bearing hemisphere; and their extent, eg. diffuse microglia activation throughout the brain.

### PVSRIPO induces acute IFN-dominant inflammation

To study PVSRIPO-induced transcriptomic landscape alterations, we selected 20 CT2A^hCD155^ specimens (**Supplementary Table 3**) collected at different intervals pre- and post PVSRIPO/mock infusion. Tumor region RNA was recovered from FFPE tissue sections for bulk RNA sequencing. Differential expression analysis revealed PVSRIPO-induced up-regulation of 380 and down-regulation of 10 genes (applying a log_2_-fold change cut-off of 2.0 and p-value cut-off of 0.05) compared to mock controls at day 8 (**Fig. 6a**). This ratio dropped to 18 and 7 genes, respectively, at day 12 (**Fig. 6a**). Consistent with the antitumor and inflammatory effects observed, PVSRIPO induced a profound but transient transcriptional response that peaked 48h post treatment. Gene set enrichment analyses (GSEA) interrogating Hallmark Genesets [29] revealed several recurring genesets in PVSRIPO-treated brains, led by ‘IFNα response’ and ‘IFNγ response’. Meanwhile, cell cycle progression genesets (‘Myc targets V1/2’, ‘G2M checkpoint’, ‘E2F targets’) were down-regulated (**Fig. 6b**). These results are consistent with the histological evidence for tumor regression and a broad, macrophage/microglia-centered innate antiviral inflammatory reaction.

**Fig. 6.**
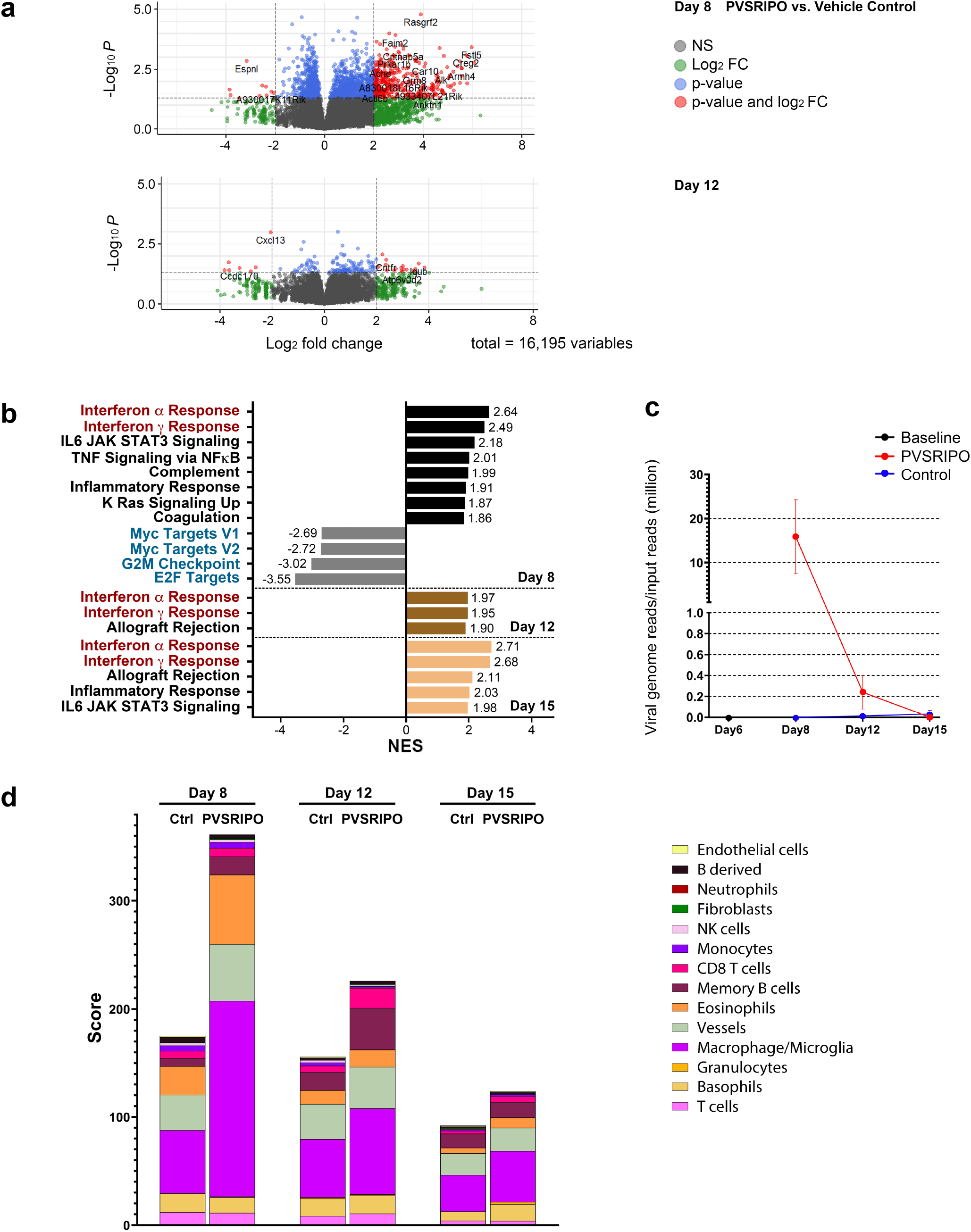
PVSRIPO treatment induces sustained intratumor type I/II IFN signaling associated with transient viral replication. RNA was extracted from FFPE slides prepared from tumor-bearing brains, roughly comprising the tumor-bearing region, collected on the indicated days post mock/PVSRIPO treatment. Extracted bulk RNA underwent rRNA depletion and library preparation, followed by sequencing (Illumina HiSeq). (**a**) Volcano plots depicting differentially expressed genes at early (day 8) and late (day 12) post-treatment intervals (a log_2_ fold change cut-off of 2.0; and an adjusted p-value cut off 0.05 were implemented). (**b**) Gene Set Enrichment Analysis revealed the list of top Hallmark genesets altered by PVSRIPO therapy over time in the CT2A^hCD155^ tumor TME, ranked in descending order of normalized enrichment score (NES). Positive NES values reflect up- and negative NES values reflect down-regulation. Only statistically significant (False Discover Rate-adjusted) genesets with NES>1.8 are shown. (**c**) Recovery of PVSRIPO viral RNA from the TME over time, represented by number of reads mapping to the poliovirus genome in each bulk RNA sample. (**d**) Computationally predicted cell type densities in the submitted samples using the MCP-counter method.[3]

RNAseq also provided an opportunity to assess viral RNA (vRNA) counts in the TME (**Fig. 6c**). There were no reads mapping to vRNA at baseline, prior to PVSRIPO infusion (day 6) or in any mock-treated samples (**Fig. 6c**). At day 8, 48h post PVSRIPO infusion, we detected 15.9-per million input vRNA reads. After day 8, the estimated viral load quickly diminished over time (**Fig. 6c**), in agreement with prior data in mouse tumor models treated with PVSRIPO [8, 10, 23]. Finally, we employed the mouse version of the Microenvironment Cell Populations (MCP)-counter algorithm [3] to calculate the abundance of tissue-infiltrating immune and stromal cells in the TME over time (**Fig. 6d**). PVSRIPO induced tumor infiltration of myeloid-derived innate immune cells at day 8 (**Fig. 6d**). This was followed by infiltration of CD8^+^T cells and memory B cells on day 12 (**Fig. 6d**). The total amount of immune/stromal infiltrates diminished over time, in step with progressive resolution of the acute inflammatory response elicited by PVSRIPO therapy (**Fig. 6d**).

### Combination of PVSRIPO with PD1:PD-L1 immune checkpoint inhibitors improves long-term antitumor efficacy in GAMM dependent manner

Robust type-I IFN-dominant antiviral inflammation with widespread PD-L1 induction against the backdrop of a profoundly engaged myeloid infiltrate, suggests that PVSRIPO’s antitumor effects could be buttressed with PD1:PD-L1 immune checkpoint blockade. Thus, we assessed antitumor efficacy of PVSRIPO combined with α-PD-L1 (**Fig. 7**) or α-PD1 (**Supplementary Fig. 7a**). To mimic the clinical scenario with pre-existing anti-PV immunity and a PV vaccine boost given prior to PVSRIPO infusion [15], mice were immunized against PV 47 and 32 days prior to CT2A^hCD155^ implantation (**Fig. 7**). CT2A^hCD155^ tumor-bearing mice were treated with mock/PVSRIPO as shown (**Fig. 1**), followed by α-PD-L1 or isotype-matched control antibodies (**Fig. 7**). Combining PVSRIPO with α-PD-L1 had a minor effect on median survival (24 days vs. 19 days for the mock + IgG group; **Fig. 7a**), but a significant effect on overall survival (OS) with 36% complete remission at day 60 (**Fig. 7a**). PVSRIPO + α-PD-L1 therapy-induced remission was resistant to contralateral rechallenge with CT2A (**Fig. 7a**), which was lethal in 6 age-matched, treatment-naïve control mice. The extension of OS with PVSRIPO + α-PD-L1 was eliminated upon CD8^+^T cell depletion (**Fig. 7b**).

**Fig. 7.**
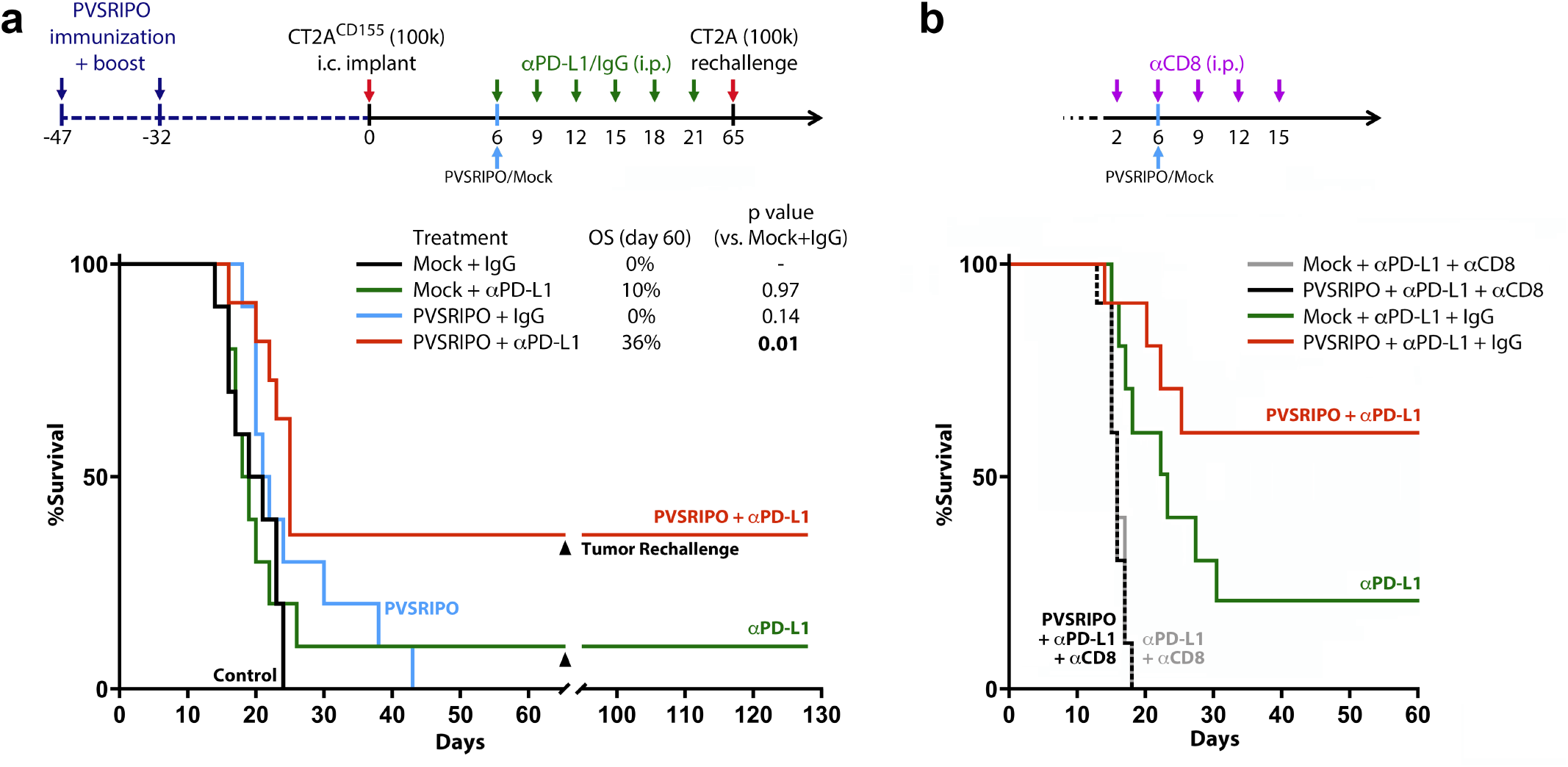
Combination therapy with PD-L1 immune checkpoint blockade enhanced long-term survival/remission in CT2A^hCD155^ glioma in a CD8^+^T cell dependent manner. (**a**) hCD155-tg mice were immunized against PV as shown (top panel), implanted with CT2A^hCD155^ glioma, and treated with PVSRIPO/mock control (DMEM) on day 6, in combination with anti-PD-L1 antibody or isotype control intraperitoneally (i.p.; 250mg per dose, in 100ml PBS) as shown. Survival for all groups (n=10; PVSRIPO + anti-PD-L1, n=11) is indicated. Mice in remission were re-challenged with 10^5^ CT2A^hCD155^ tumor cells in the contralateral brain (day 65), in parallel with age-matched treatment-naive mice as controls (n=5). The p-values shown are from Mantel-Cox log-rank test, two tailed. (**b**) Adding CD8^+^T cell depletion (with i.p. anti-CD8 antibody as shown) to the study regimen shown in (a) eliminated the PVSRIPO + anti-PD-L1 and anti-PD-L1 treatment effects.

To assess the role of PVSRIPO-induced GAMM engagement in immunotherapy efficacy of PVSRIPO + α-PD-L1 and of α-PD-L1 alone, treatment was repeated in animals treated with clodronate liposome-mediated macrophage depletion (**Supplementary Fig. 7b**). Incomplete GAMM depletion (∼30% reduction of GAMM per CD68^+^ staining) decreased the combination therapy effect from 40.0% to 16.7% (**Supplementary Fig. 7b**).

## Discussion

From their inception [31], viral anti-glioma strategies were predicated on a concept of viral infection and cytopathogenicity in neoplastic cells, based on robust virus growth in high-passage *in vitro* tumor models. In this context, CNS resident innate immune cells often were deemed an obstacle to therapy due to their role in clearing viral agents administered intratumorally [18].

PVSRIPO, by virtue of widespread CD155 upregulation in malignancy [24], including malignant glioma [12, 33], infects and damages cancerous cells via antiviral IFN responses [46] and cytotoxic viral proteins *in vitro* [6, 7]. Yet, studies of PVSRIPO in *ex vivo* GBM slices, a clinically relevant model with authentic cellular heterogeneity, show that neoplastic cells are not the main target for its biological activities [10]. Our studies in I.C. mouse tumor models confirm that PVSRIPO mainly provokes responses in the GAMM compartment *in vivo*, unleashing type-I IFN dominant engagement of the notoriously immunosuppressive glioma myeloid infiltrate. This reverberates with PV’s natural cell type-specificity for migratory/lymphoid DCs and macrophages [44].

PVSRIPO therapy of I.C. mouse tumor models caused a rapid decrease of tumor size evident as reduced Ki67^+^ cells in the tumor bed, discohesive architecture, and a decline in cell cycle- and proliferation-related gene expression. Absent vRNA accumulation in the tumor all but excludes the possibility of direct viral cell killing to account for this. This is consistent with PVSRIPO therapy of hCD155-tg hosts (implying PVSRIPO targeting of myeloid cells) occurring with hCD155-negative tumor implants that are resistant to infection [10]. Indeed, CD8^+^T cell depletion and clodronate liposome studies indicate that PVSRIPO-instigated antitumor effects are immune mediated. Yet, we cannot rule out a role for viral infection of- and cytopathogenic damage to malignant cells in the antitumor effect, even if it is sub-lethal and non-productive.

The main histological finding after PVSRIPO was marked microglia activation engulfing virtually the entire brain. Microglia are the bulwark of the CNS innate antiviral response [14]. ‘Classic’ viral encephalitis, eg. the syndromes modeled in mice infected with rhabdo- (vesicular stomatitis virus) [13], flavi- (eg. West Nile virus) [42], or neurotropic coronaviruses [26], share common features implicated in triggering microglia activation [14]. They each infect neurons and inflict substantial neuronal damage; they infect and damage astro- and/or oligodendrocytes; they induce peripheral monocytic infiltration in the CNS. In viral encephalitis models, microglia activation occurs in close proximity to these triggers, eg. foci with viral propagation in- and damage to CNS cells [37]. PVSRIPO CNS infection lacks all of these triggers. PVSRIPO is devoid of cytopathogenic potential in neurons [34]; it is, thus, non-neuropathogenic in human subjects [15], non-human primates [16], and hCD155-tg mice [22]. PVs do not infect astro-/oligodendrocytes. I.C. PVSRIPO did not induce peripheral monocyte infiltration beyond the needle track.

Unambiguously identifying microglia in the virus-infected CNS is difficult, especially against the backdrop of peripheral monocytic infiltration [14]. Tmem119 staining of cells with morphological features of activated microglia defines this compartment to respond to PVSRIPO. There was profuse proliferation of such cells, evident as Ki67^+^ co-staining, in the brain. Microglia activation and proliferation induced by PVSRIPO, unlike classic viral encephalitis, was diffuse and extensive, consistent with the absence of focal viral cytopathogenicity. Rather, this pattern of microglia activation may reflect PVSRIPO’s non-lethal phenotype in cells of the mononuclear phagocytic system yielding sustained type-I IFN dominant inflammation [9, 10, 36]. Accordingly, the prevailing transcriptional signatures emerging from RNAseq analyses of the tumor region post PVSRIPO were antiviral inflammatory in nature, including upregulation of PD-L1.

The dramatic early treatment effects of PVSRIPO were transient and, in ∼90% of cases, gave rise to lethal tumor progression. Our studies suggest that turning acute inflammatory antiviral reactions into durable remission may be achieved by combining PVSRIPO with immunomodulatory agents capable of blocking natural resistance points of the antiviral immune response, eg. with anti-PD-L1 or PD1 immune checkpoint blockade.

Our investigations demonstrate that, rather than a detriment to cancer virotherapy, the GAMM infiltrate is a promising substrate for viral immune stimulatory actions. Beyond proinflammatory engagement of the tumor myeloid infiltrate, the quality, depth, and anatomical reach of PVSRIPO-induced microglia activation may be indispensable for generating functional antitumor CD8^+^T cell immunity in malignant gliomas, notorious immune deserts intrinsically resistant to systemic immunotherapy approaches [20].

## Acknowledgement

We thank Dr. Z. Su and H. Dai for technical assistance with IHC, Dr. E. Dobrikova for technical assistance with immunoblots, P. Healy for statistical assistance, Dr. S. Koike (Tokyo Metropolitan Institute of Medical Science, Japan) for providing the hCD155-tg C57Bl6 mice, Dr. S. Keir for animal training, and Drs. J. Bryant and, M. Mosaheb for assistance with animal surgeries. We thank Dr. R.E. McLendon for critical reading of the manuscript.

