## Supplemental Material for "Polio Virotherapy of Malignant Glioma Engages the Tumor Myeloid Infiltrate and Induces Diffuse Microglia Activation"

Yuanfan Yang *et al*.

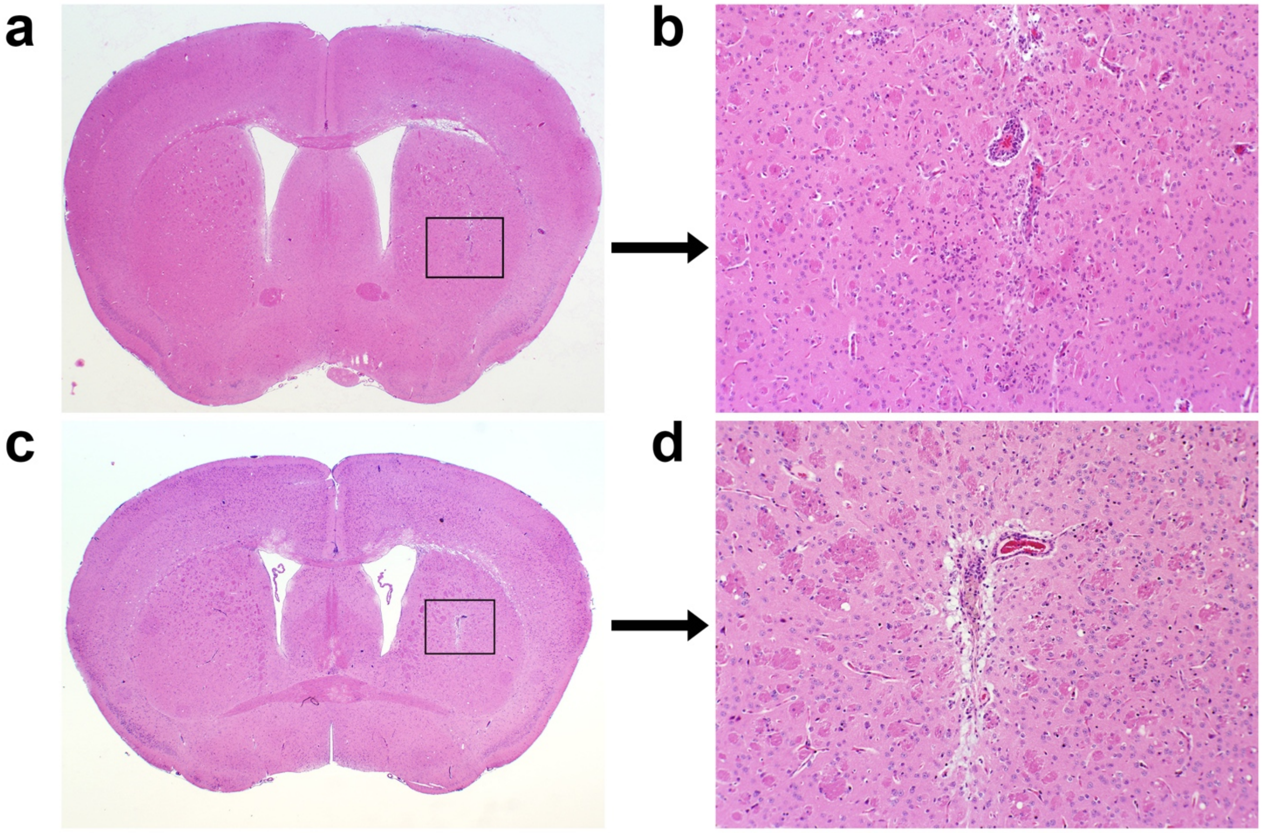


**Supplementary Fig. 1** (related to Fig. 1). H&E histology of coronal FFPE sections from mice in pathological remission at day 15 (**a**, **b**) or day 18 (**c**, **d**) post PVSRIPO. Sections are from cases nr. 80 (**a**, **b**) or R16 (**c**, **d**; see **Supplementary Table 3**)**.** The approximate locations of the tumor implantation sites are indicated by black boxes (**a**, **c**).

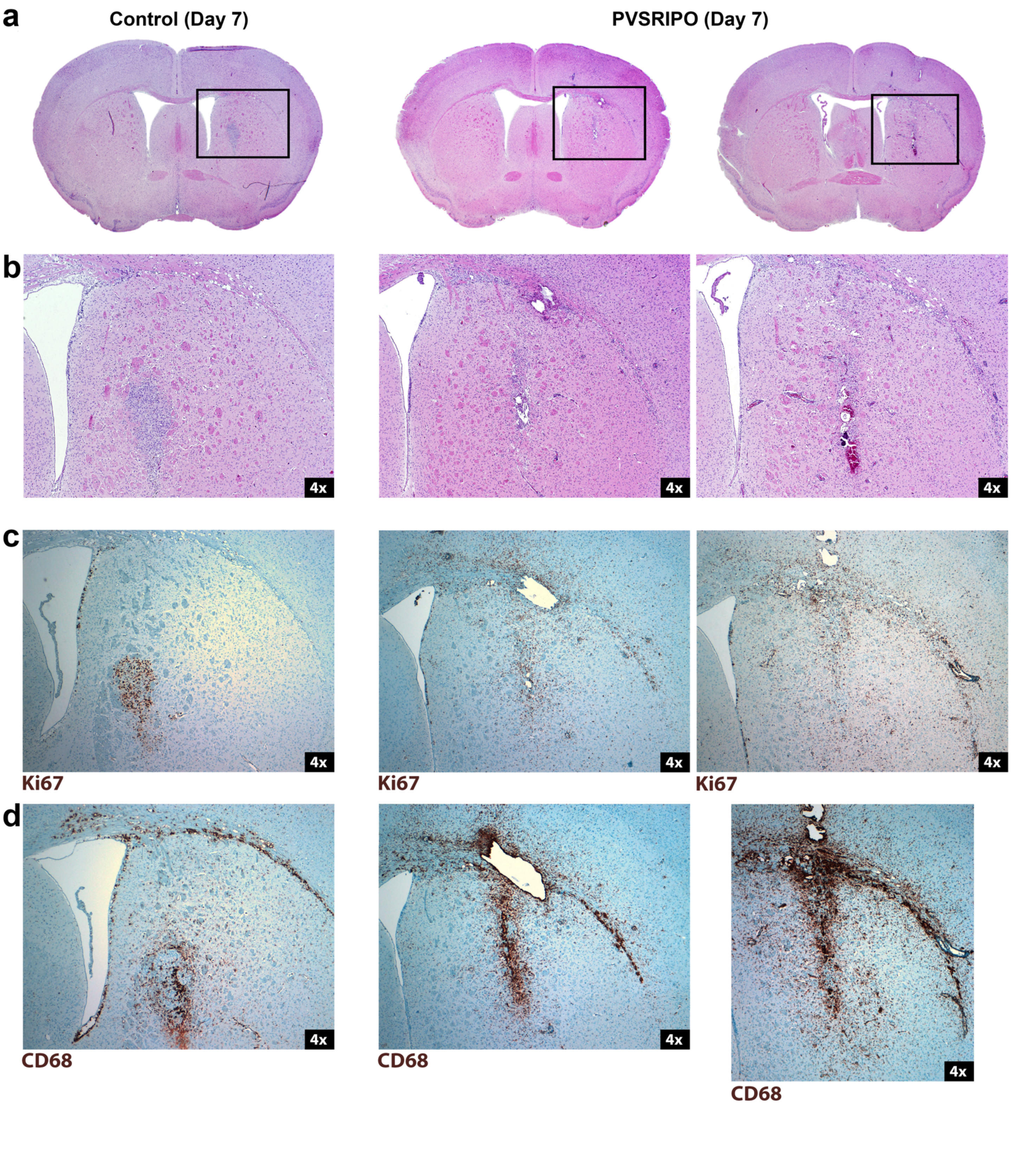
a
Control (Day 7)
PVSRIPO (Day 7)
b
4x
4x
4x
c
4x
4x
4x
Ki67
Ki67
Ki67
d
4x
4x
4x
CD68
CD68
CD68

**Supplementary Fig. 2** (related to Fig. 2). The treatment response to PVSRIPO in an I.C. $B16^{hCD155}$ mouse melanoma model. (**a**, **b**) H&E histology of $B16^{hCD155}$ treated with mock (left panels) or PVSRIPO (middle and right panels) at day 7 post implantation, 48h post intratumor treatment. H&E histology of the brain (**a**) and the tumor implantation site at higher magnification (**b**). Tumor regress and discohesion were evident as disorganized, reduced Ki67 staining at the tumor implantation site of PVSRIPO-treated mice compared to mock-treated controls (**c**). Profuse neuroinflammatory reactions upon PVSRIPO therapy were evident as markedly increased signal with CD68 staining (**d**).

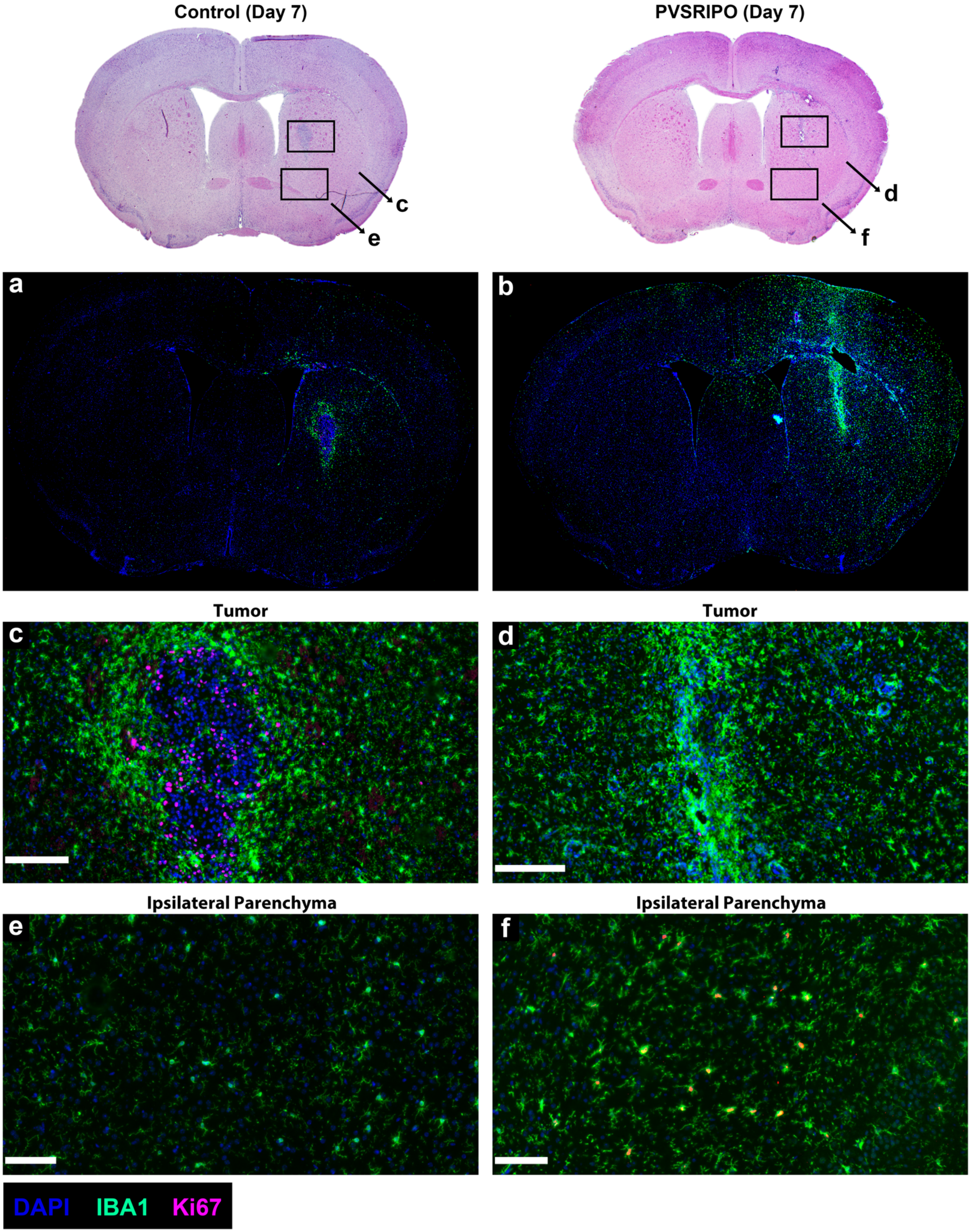
Control (Day 7)
PVSRIPO (Day 7)
c
e
d
f
a
b
Tumor
Tumor
c
d
Ipsilateral Parenchyma
Ipsilateral Parenchyma
e
f
DAPI
IBA1
Ki67

**Supplementary Fig. 3** (related to Fig. 4). IF analyses of the PVSRIPO treatment response in the B16$^{hCD155}$ model. The H&E panels atop represent mock- (left) and PVSRIPO-treated (right) brains at day 7 (**Supplementary Fig. 2a**). (**a**, **b**) Iba1 staining revealed the extent of GAMM activation in the tumor bed and microglia engagement engulfing ipsi- and contralateral hemispheres. (**c**, **d**) Co-staining of Iba1 with Ki67 documents the distribution of Iba1$^{+}$ staining GAMM in relation to the tumor (**c**), or the tumor bed after tumor regression (**d**). Size bars = 200 μm. (**e**, **f**) Co-staining of Iba1 with Ki67 revealed evidence for microglia proliferation in parenchyma away from tumor in PVSRIPO-treated animals only. Size bars = 100 μm.

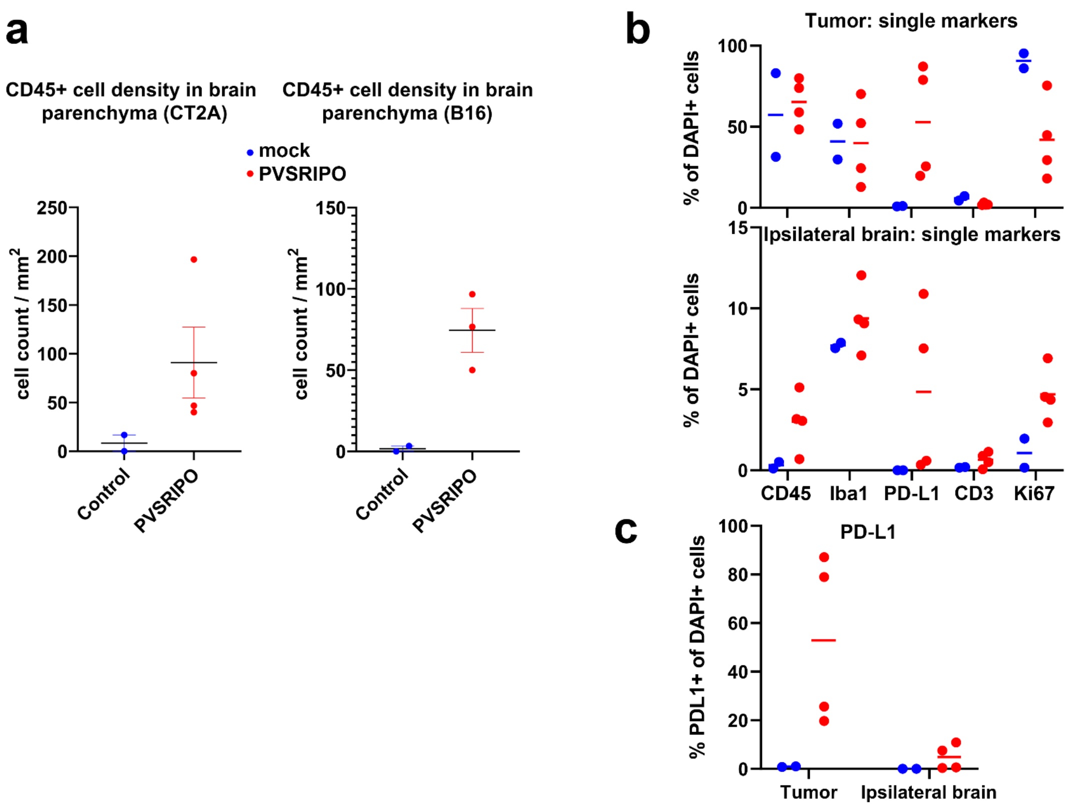


**Supplementary Fig. 4** (related to Figs. 4, 5). Quantification of various marker-positive cell populations (see text for detail). (**a**) Tissue sections from CT2A$^{hCD155}$ and B16$^{hCD155}$ bearing mice stained for CD45$^+$ IHC were evaluated for the number of positive cells by visual inspection. The CD45$^+$ cell count per mm$^2$ in mock vs. PVSRIPO treated animals is shown. (**b**) Automated, computer-based (IF) signal quantification of marker-positive staining cells expressing CD45, Iba1, PD-L1, CD3 or Ki67 in the tumor area (top) or ipsilateral normal brain (bottom). (**c**) Automated, computer-based (IF) signal quantification of PD-L1 positive cells in the tumor area and in ipsilateral normal brain shown on the same scale.

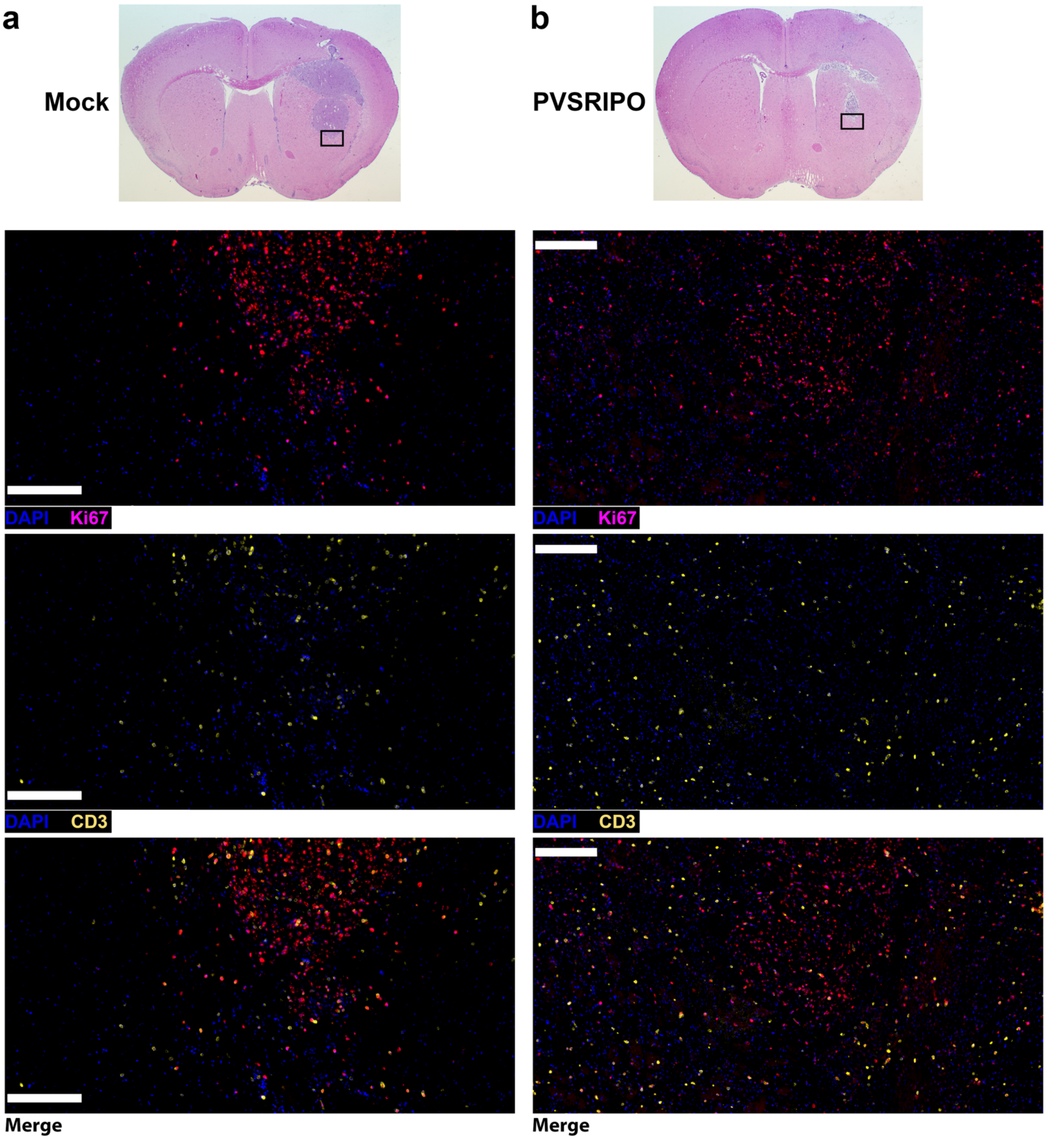
a
Mock
b
PVSRIPO
DAPI Ki67
DAPI CD3
Merge
DAPI Ki67
DAPI CD3
Merge

**Supplementary Fig. 5** (related to Figs. 4, 5). PVSRIPO therapy in the $CT2A^{hCD155}$ model induces $CD3^+$ cell infiltration and proliferation in the tumor periphery. Mock- (**a**) and PVSRIPO-treated (**b**) $CT2A^{hCD155}$ tumors (day 8) were stained for Ki67 and CD3 expression as shown. Scale bars = 100 μm.

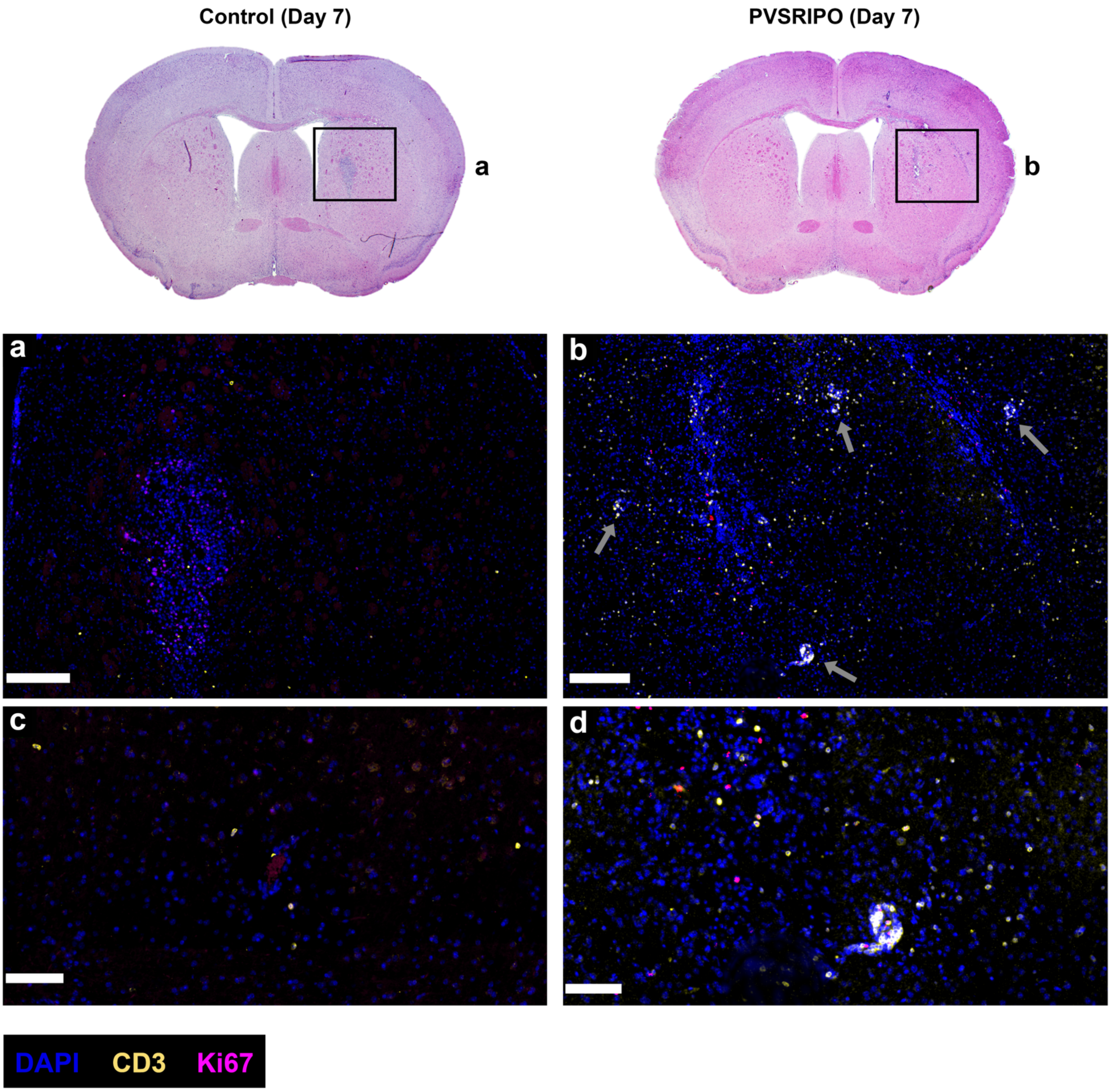
Control (Day 7)
PVSRIPO (Day 7)
a
b
a
b
c
d
DAPI
CD3
Ki67

**Supplementary Fig. 6** (related to Figs. 4, 5). IF analyses of the PVSRIPO treatment response in the $B16^{hCD155}$ model. The H&E panels atop represent mock- (left) and PVSRIPO-treated brains at day 7 (**Supplementary Fig. 2a**). (**a**-**d**) Merged staining of the tumor implantation site in overview (**a**, **b**; size bars = 200 μm) and adjacent parenchyma showing perivascular inflammation in detail (**c, d**; size bars = 200 μm); arrows point to peritumoral $CD45^+$ infiltration/perivascular cuffs in PVSRIPO-treated animals.

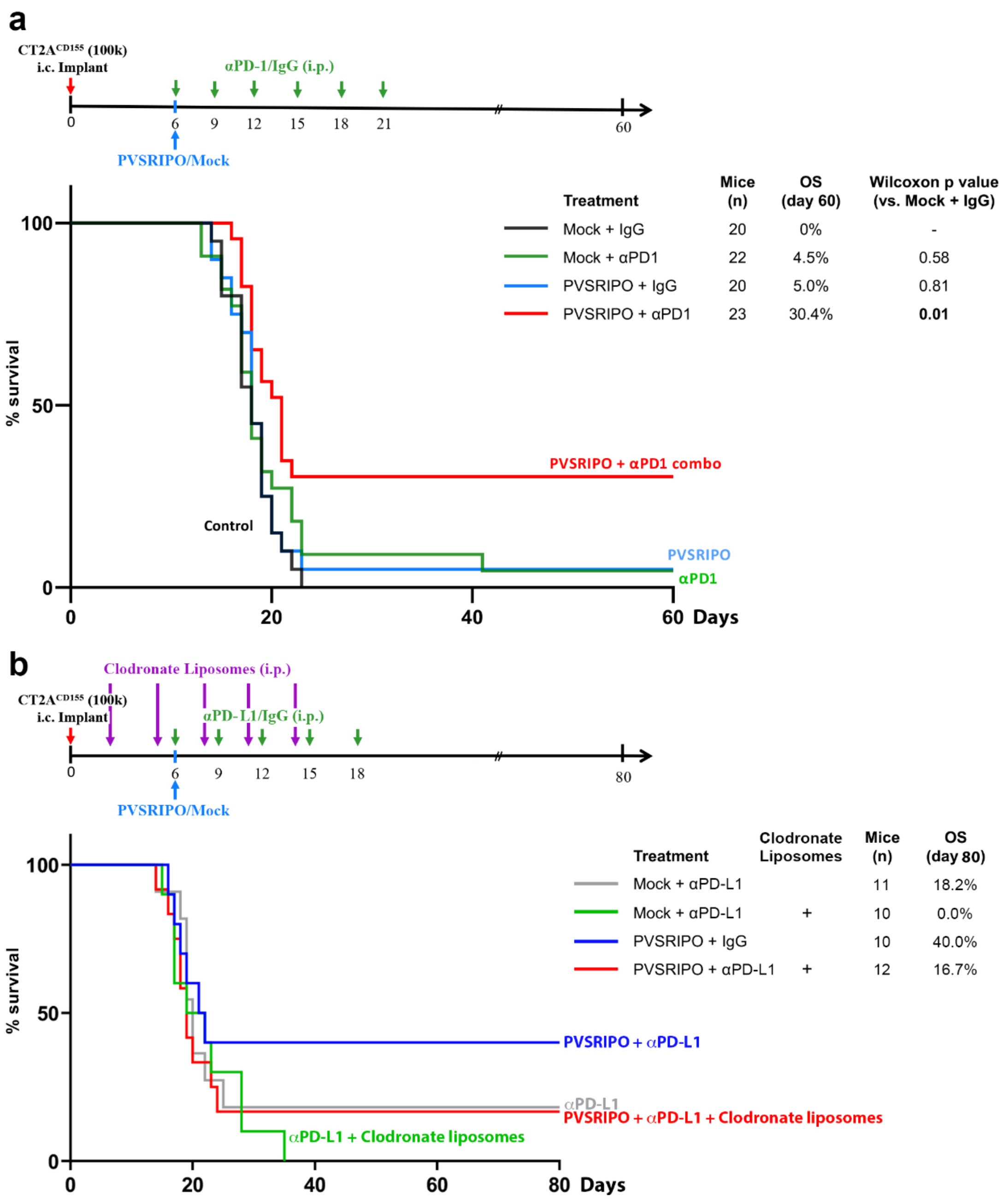


**Supplementary Fig. 7** (related to Fig. 7). (**a**) Combination with PD1 immune checkpoint blockade enhanced long-term survival/remission in CT2A$^{hCD155}$ glioma upon PVSRIPO treatment. (**b**) Macrophage depletion upon i.p. administration of clodronate liposomes partially eliminated PVSRIPO + αPD-L1 therapy effects.

**Supplementary Table 1.** Key Resources.

| REAGENT or RESOURCE | SOURCE | IDENTIFIER (USE) |
|---|---|---|
| Anti-mouse PD-L1 | BioXCell | BE0101; RRID: AB 10949073 (animal study) |
| Anti-mouse PD-1 | BioXCell | BE0146; RRID: AB 10949053 (animal study) |
| Anti-mouse CD4 | BioXCell | BE0003-1; RRID: AB 1107636 (animal study) |
| Anti-mouse CD8α | BioXCell | BE0061; RRID: AB 1125541 (animal study) |
| *InVivo*MAb Rat IgG2b | BioXCell | BE0090; RRID: AB 1107780 (animal study) |
| *InVivo*MAb Rat IgG2a | BioXCell | BE0089; RRID: AB 1107769 (animal study) |
| **Anti-mouse IHC antibodies** | | |
| Anti-CD3 | Abcam | ab5690; RRID: AB 305055 (IHC) |
| Anti-CD45 | Cell Signaling | 70257; RRID: AB 2799780 (IHC) |
| Anti-Tmem119 | Cell Signaling | 80821; RRID: n/a (IHC) |
| Anti-Iba-1 | Cell Signaling | 17198; RRID: AB 2820254 (IHC) |
| Anti-CD68 | Cell Signaling | 76437; RRID: AB 2799882 (IHC) |
| Anti-Ki67 | Cell Signaling | 12202; RRID: AB 2620142 (IHC) |
| Anti-Iba1 | Novus | NB100-1028; RRID: AB 521594 (IF) |
| Anti-CD68 | Cell Signaling | 76437; RRID: AB 2799882 (IF) |
| Anti-Ki67, Alexa Fluor®647 conj. | Cell Signaling | 12075; RRID: AB 2728830 (IF) |
| Anti-PD-L1 | Novus | 76769; RRID: AB 11024101 (IF) |
| Anti-STAT1 | Cell Signaling | 9172S; RRID: AB 2198300 (immunoblot) |
| Anti-tubulin | Cell Signaling | 2148; RRID: AB 2288042 (immunoblot) |
| Anti-hCD155 | Cell Signaling | 81254; RRID: AB 2799970 (immunoblot) |
| Anti-(poliovirus) 2C | Lab-generated | N/A (immunoblot) |
| **Secondary antibodies** | | |
| DISC. OmniMap anti-Rb HRP | Roche Ventana | 760-4311; RRID: AB 2811043 (IHC) |
| Donkey α-Goat AlexaFluor®488 | Invitrogen | A-11055; RRID: AB 2534102 (IF, Iba1) |
| Donkey α-Rabbit AlexaFluor®555 | Invitrogen | A-31572; RRID: AB 162543 (IF, CD68/PD-L1) |
| **Critical Kits** | | |
| RNeasy Plus Micro Kit (50) | Qiagen | Cat#74034 (RNAseq) |

**Supplementary Table 2.** Optimizing the CT2A$^{hCD155}$ model for containment within the hemisphere receiving the implant.

| Depth (mm) | Needle (Gy) | Drill | % of contained tumor | tumor take % | No. of Animals |
|---|---|---|---|---|---|
| 2.5 | 25 | N | **27%** | 100% | 15 |
| 3.0 | 25 | N | **35%** | 100% | 20 |
| 3.0 | 27 | N | **55%** | 100% | 40 |
| 3.0 | 27 | Y | **78%** | 100% | 9 |
| 3.6 | 27 | Y | **78%** | 100% | 9 |
| 3.6 | 30 | Y | **93.3%** | 100% | 30 |

**Supplementary Table 3.** Listing of all mice implanted with CT2A$^{hCD155}$ enrolled in the study, which were used for brain harvest and histopathological/IHC analyses of PVSRIPO treatment effects.

| Study | Sample | Treatment | Study Day |
|---|---|---|---|
| 1 | 8 | - | 6 |
| 1 | 24 | - | 6 |
| 2 | R26 | - | 6 |
| 3 ■ | 51 | - | 6 |
| 2 | R27 | - | 6 |
| 3 ■ | 50 | - | 6 |
| 1 | 3 | Mock | 8 |
| 3 ■ | 54 | Mock | 8 |
| 3 ■ | 55 | Mock | 8 |
| 2 | R51 | Mock | 8 |
| 1 | 2 | PVS | 8 |
| 1 | 1 | PVS | 8 |
| 3 ■ | 57 | PVS | 8 |
| 3 ■ | 53 | PVS | 8 |
| 3 ■ | 56 | PVS | 8 |
| 3 ■ | 52 | PVS | 8 |
| 2 | R52 | PVS | 8 |
| 2 | R50 | PVS | 8 |
| 1 | 22 | Mock | 10 |
| 1 | 21 | Mock | 10 |
| 3 | 61 | Mock | 10 |
| 3 ■ | 60 | Mock | 10 |
| 3 | 58 | PVS | 10 |
| 1 | 23 | PVS | 10 |
| 3 | 62 | PVS | 10 |
| 3 | 63 | PVS | 10 |
| 3 | 59 | PVS | 10 |
| 3 ■ | 68 | Mock | 12 |
| 3 ■ | 69 | Mock | 12 |
| 2 | R48 | Mock | 12 |
| 2 | R45 | Mock | 12 |
| 3 | 83 | PVS | 12 |
| 3 ■ | 65 | PVS | 12 |
| 3 ■ | 64 | PVS | 12 |
| 3 ■ | 66 | PVS | 12 |
| 3 ■ | 67 | PVS | 12 |
| 2 | R44 | PVS | 12 |
| 2 | R46 | PVS | 12 |
| 3 ■ | 74 | Mock | 15 |
| 3 ■ | 76 | Mock | 15 |
| 2 | R32 | Mock | 15 |
| 2 | R34 | Mock | 15 |
| 3 ■ | 72 | PVS | 15 |
| 3 ■ | 71 | PVS | 15 |
| 3 ■ | 73 | PVS | 15 |
| 3 | 80 | PVS | 15 |
| 2 | R35 | PVS | 15 |
| 2 | R33 | PVS | 15 |
| 2 | R43 | PVS | 15 |
| 2 | R36 | Mock | 18 |
| 1 | 20 | Mock | 18 |
| 1 | 18 | PVS | 18 |
| 1 | 19 | PVS | 18 |
| 2 | R31 | PVS | 18 |
| 2 | R16 | PVS | 18 |

Brains from these 49 animals were FFPE processed and submitted for blinded histopathological analysis

■ FFPE sections from the brains of these 20 animals were submitted for RNAseq analysis

Brains from these 6 animals were FFPE processed, but not included with the other 49 samples for analyses, due to sample attrition
